# Probabilistic Ranking Of Microbiomes Plus Taxa Selection to discover and validate microbiome function models for multiple litter decomposition studies

**DOI:** 10.1101/2020.07.17.209031

**Authors:** Jaron Thompson, Nicholas Lubbers, Marie E. Kroeger, Rae DeVan, Renee Johansen, John Dunbar, Brian Munsky

## Abstract

The overwhelming complexity of microbiomes makes it difficult to decipher functional relationships between specific microbes and ecosystem properties. While machine learning analyses have demonstrated an impressive ability to correlate microbial community composition with macroscopic functions, mechanisms that dictate model predictions are often unknown, and predictions often lack an assigned metric of uncertainty. In this study, we apply Bayesian networks to build on prior feature selection analyses and construct easy-to-interpret probabilistic models, which accurately predict levels of dissolved organic carbon (DOC) from the relative abundance of soil bacteria (16S rRNA gene profiles). In addition to standard cross-validation, we show that a Bayesian network model trained using samples from a pine litter decomposition study accurately predicts DOC of samples from an independent oak litter decomposition study, suggesting that mechanisms driving variation in soil carbon storage may be conserved across different types of decomposing plant litter. Furthermore, the structure of the resulting Bayesian network model defines a minimal set of highly informative taxa, whose abundances directly constrain the probability of high or low DOC conditions. Significant accuracy of the Bayesian network model with independent data sets supports the validity of the identified relationships between taxa abundance and DOC.

**Summary:** Understanding the interplay between microbiomes and the environments they inhabit is a daunting task. While recent advances in gene sequencing technology provide a means of profiling the relative abundance of microbial species, the resulting data are noisy, sparse, and limited to small sample sizes. Despite these challenges, machine learning approaches have demonstrated a promising ability to discover patterns linking the microbiome with macroscopic behavior. However, most machine learning models applied to microbiome data do not estimate prediction uncertainty and provide little insight regarding how predictions are made. In this study, we couple machine learning approaches for feature reduction with Bayesian networks to model the relationship between the soil microbiome and dissolved organic carbon (DOC). We show that Bayesian networks are accurate and provide a transparent link between microbial abundance and DOC. To validate Bayesian networks, we demonstrate accurate predictions for held-out testing data and with data from independent decomposition experiments.

## Introduction

Microbiome functions are extremely diverse and can potentially be optimized to efficiently perform industrially relevant tasks. However, engineering microbiomes requires validated models that accurately predict functions from microbial community profiles. Analytical techniques to infer microbial taxa that control macroscopic functions must overcome several obstacles. First, microbiomes are often composed of thousands of bacterial and fungal species [1]. Second, measuring microbiome composition by DNA sequencing has large measurement uncertainty, and studies are often limited to a small collection of experimental observations. Third, microbiomes are typically spatially heterogeneous, which increases measurement noise. Fourth, the composition of microbiomes varies by ecosystem [2], creating additional compositional variations that impede efforts to identify “universal” taxa that drive community functions.

Machine learning approaches are well-suited to deal with these complexities, with proven success in reducing the large microbiome feature spaces to small subsets of microbial taxa that are most informative for prediction tasks [3]. Machine learning techniques, such as random forest algorithms, have demonstrated impressive prediction accuracy with microbiome data [4, 5]. However, many machine learning workflows often suffer from the following limitations: First, machine learning models that are typically applied to metagenomic data lack the ability to quantify prediction uncertainty, which prevents users from knowing whether or not a prediction should be trusted in a new experimental circumstance. Second, machine learning analyses of microbiome data are often considered to be a “black box”, in which the mechanisms that underpin model predictions are unknown or difficult to conceptualize, which makes models less useful from an engineering perspective. Finally, machine learning models are rarely validated with data from independent studies, relying only on cross-validation of held-out testing data to confirm model generalizability [6].

In regard to uncertainty analysis and model interpretation, probabilistic machine learning models offer clear advantages over deterministic models. Probabilistic models provide quantified prediction confidence that reflect uncertainty resulting from noise in the data and limited sample sizes [7]. One particular probabilistic modeling framework that has gained popularity in modeling biological processes and elsewhere is the *probabilistic graphical model* [8–10]. PGMs not only provide estimates of prediction uncertainty, but the graphical structure of PGMs addresses the issue of model interpretability by providing a visualizable interpretation of how variables influence model predictions. Probabilistic models, such as PGMs, can also be more insightful than deterministic models in cross-study validation analyses, in that PGM prediction uncertainty is amplified when extrapolated to poorly constrained circumstances, thus enabling users to gauge model relevance for new testing experiments.

A *Bayesian network* is a specific type of PGM that connects model variables with directed edges to describe the dependence properties of model variables [11]. The directed graph structure of Bayesian networks allows the user to identify a *minimum* set of features, known as a Markov blanket (see methods) that directly constrains the probability of the target variable or variables that the user wishes to predict. The combination of probabilistic predictions, model interpretability, and feature reduction make Bayesian networks an attractive approach for modeling microbial community behavior. However, beyond a few pilot studies [12, 13], the use of Bayesian networks for modeling microbial community behavior remains rare [14] due to the computational cost of learning Bayesian networks over many variables.

In this article, we apply Bayesian networks to predict levels of dissolved organic carbon (DOC) from the abundances of microbial taxa in decomposing plant litter. This case study is motivated by the need to understand the role of the soil microbiome in carbon cycling. The accumulation of greenhouse gases, especially carbon dioxide (CO_2_), in the atmosphere is a primary cause of global warming [15]. One potential mitigation strategy to offset greenhouse gas emissions is to increase soil carbon sequestration, which has the potential to reduce emissions by 0.4 to 1.2 gigatons of carbon per year [16]. In terrestrial ecosystems, the fate of carbon (i.e., whether released as CO_2_ or stored as soil organic matter) is largely determined by microorganisms [17]. As microorganisms metabolize plant litter, CO_2_ is released from microbial respiration, and a variety of dissolved carbon compounds (e.g., sugars, peptides, lipids) that may be stabilized as soil organic matter are released either from deconstructed plant material or from microbial cells [2,18]. While the composition of the soil microbiome is known to significantly influence the fate of carbon in soil, methods to elucidate specific microbial taxa that drive carbon fate are limited [19]. Overcoming this methodological challenge could enable the design of microbial communities that promote increased soil carbon storage. More powerful data analysis techniques may be a solution.

To address the high dimensionality of microbiome data, we first apply Random Forest, Indicator species and Neural Network (RFINN) feature-reduction framework [3] to determine the features (i.e., microbial taxa) that are most informative to predict DOC. We then train a Bayesian network structure over this reduced set of features to determine a set of bacterial genera whose abundances directly influence the probability of high or low DOC. We then use cross-validation to verify that the model is predictive for held out testing data, and we use cross-study validation to investigate model generalizability and transferability. Our results suggest that the mechanisms that drive variation in soil carbon storage may be conserved at least partially across litter types. Furthermore, we show that trained Bayesian networks provide reliable estimates of prediction fidelity, where more certain predictions are generally more accurate than less certain predictions. Ultimately, high-confidence predictions of such models could be used to suggest novel communities that are more likely to promote accumulation of soil organic carbon.

## Results

Microbial abundance and dissolved organic carbon (DOC) data were collected from plant litter decomposition experiments performed in laboratory microcosms [5,20]. Microcosms containing sterilized plant litter were inoculated with microbial community samples taken from 206 different locations throughout the southwestern United States and left to incubate for a period of 44 days. After the incubation period, DOC and microbial community abundances (16S rRNA gene profiles) were recorded. For all modeling analyses in this study, relative abundance of bacterial genera was used as features for prediction of DOC. For a more detailed account of litter decomposition experiments and data pre-processing, see materials and methods.

From the measured bacterial genera abundances, we created a discretized data matrix,

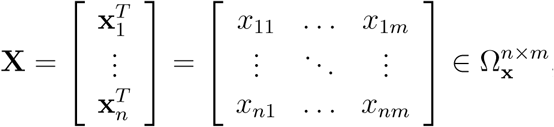

to represent the abundances of *m* microbial genera in each of the *n* different training samples, where each *x*_*ij*_ ∈ Ω_**x**_ = {Low, Moderate, High}. Similarly, we let 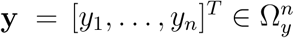 denote the discretized abundance of DOC for the same samples, where *y*_*i*_ ∈ Ω_*y*_ = {Low, High}. Bin sizes for features and targets, Ω_**x**_ and Ω_*y*_, were defined based solely on the training data.

Figure 1 provides a general overview of our approach, which we call PROMPTS (Probabilistic Ranking Of Microbiomes Plus Taxa Selection), to learn and validate a probabilistic model to predict environmental traits from relative microbial abundances. Specifically, our goal is to use PROMPTS to construct a model that estimates DOC levels (*y*) from microbial composition (**x**). In the next subsection, we analyze pine litter data in the PROMPTS framework to constrain a simple Bayesian network that models the joint probability distribution of the DOC level and genera abundances (i.e., *P* (*y*, **x**)). We then use five-fold cross-validation to verify that the estimated conditional probability of DOC given observed genera abundance (i.e., *P* (*y* | **x**)) provides accurate estimates of DOC levels from held-out data from pine litter experiments. Finally, we use cross-study validation to test how well the PROMPTS analysis is able to predict DOC variation for new oak and grass litter experiments.

**Fig 1.**
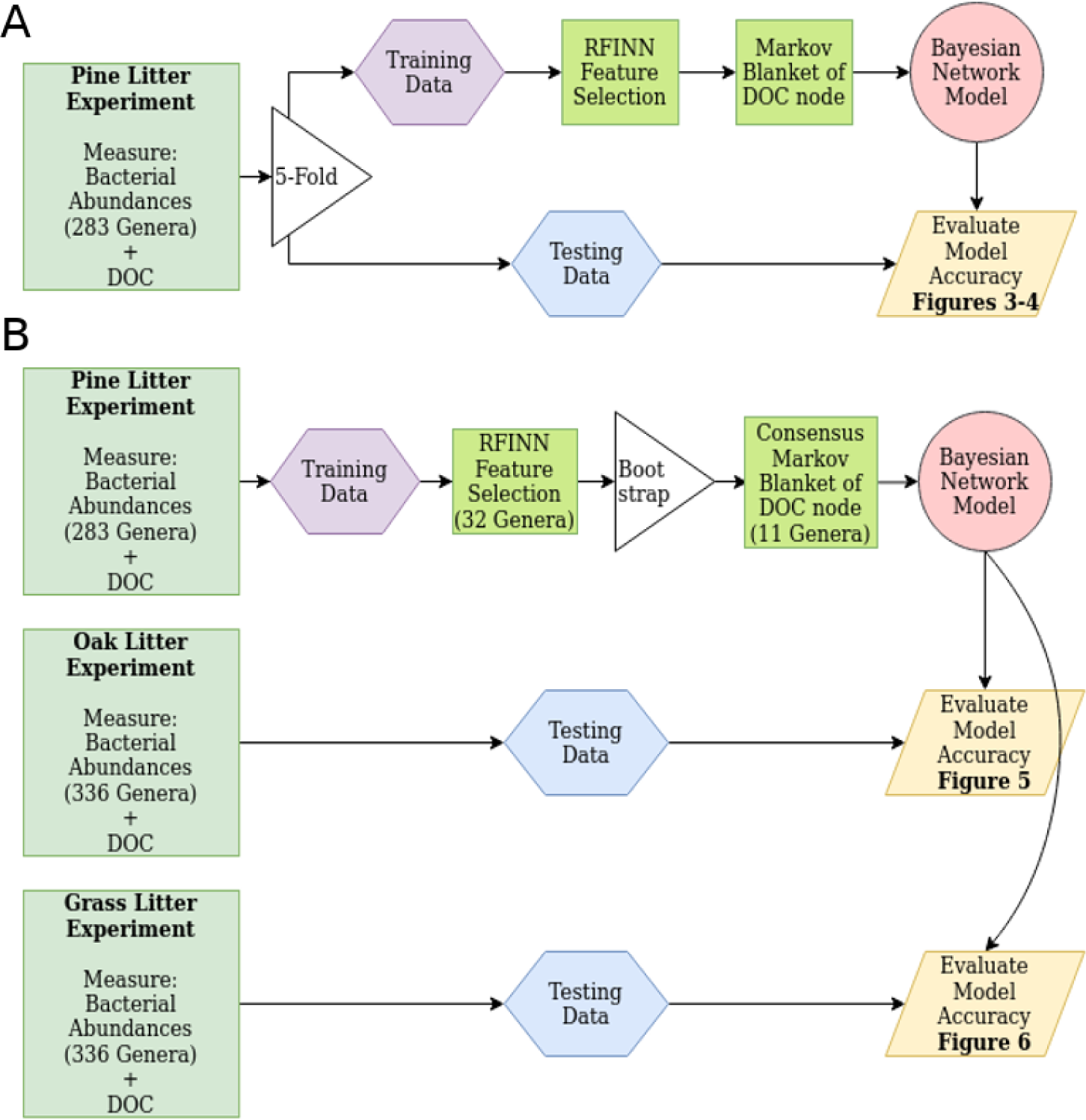
Schematic of the PROMPTS approach for feature selection, training, and validation strategies. (A) Training and validation with pine litter data: The full data set is partitioned into five permutations of training and held-out testing data. RFINN is applied to each permutation of training data to reduce the feature set. Then, the grow-shrink algorithm is used to find the Markov blanket of the DOC node and train a local BN. Prediction accuracy of the BN model is validated on held-out testing data. (B) Training and validation with validation data from independent litter decomposition experiments: For cross-study validation, RFINN is first applied to the entire pine data set for feature reduction. Then, for 30 random data sub-samples, a grow-shrink algorithm is performed to identify DOC’s Markov blanket and train a local BN. The prediction accuracy of the consensus BN is validated against oak and grass data.

### Key relationships between microbiome abundances and functional outputs can be captured by a small and robust Bayesian Network

Using training data (**X, y**) as described above, PROMPTS seeks to constrain a probabilistic graphical model (PGM) to estimate *P* (*y* | **x**) and represent the associations between microbial genera (**x**) and DOC (*y*). In this PGM representation, a graph, 𝒢, is composed of a set of nodes, 𝒩, that represent random variables and edges, ℰ, that represent the probabilistic dependence between nodes. In our case, PROMPTS focuses on a specific type of PGM known as a Bayesian network (BN), where each of *m* feature nodes describes the abundance of a specific genus; one target node describes the level of DOC; and the graphical structure imposes conditional independence properties between nodes [7, 11].

Any joint distribution *P* (*y*, **x**) could be represented by a fully connected graph in which all node pairs are connected by an edge. However, conditional independences between nodes allow for the removal of edges to form graphs that are simpler, but equally valid, representations of *P* (*y*, **x**) [7]. In practice, determining the structure of a BN without prior knowledge requires the completion of a structure learning search algorithm. One approach to determine the optimal Bayesian network structure is to search the space of possible graph structures and return the configuration with the best score. A common scoring metric is the *minimum descriptive length*, which encourages model fitness and penalizes model complexity (i.e. the number of edges) [21]. Unfortunately, the computational effort for structure learning of BNs scales super-exponentially in the number of variables [11], which makes exhaustive structure searches highly impractical for microbiome data that contain hundreds of taxonomic features. For this reason, PROMPTS first applies an integrated framework of Random Forest, Indicator species, and Neural Network algorithms (RFINN [3]) to down-select from the original data containing 283 genera to a consensus set of 32 genera that were significant predictors of DOC for all three algorithms.

Even after feature reduction by RFINN, an exhaustive structure search over the reduced set of 32 variables to find a BN to estimate *P* (*y*, **x**) would remain computationally expensive [22]. However, our specific goal for this study is to estimate *P* (*y* | **x**) to quantify how knowledge of microbial features constrains uncertainty in the level of DOC. In addition to reducing the required number of edges in a BN, conditional independences can also substantially reduce the number of nodes needed to constrain the conditional probability of a target variable. The *Markov blanket* for any target node, *y*, is defined as the minimum set of nodes that, when observed, render *y* conditionally independent of all other nodes in the graph (e.g., the minimal set of genera that are needed to predict the level of DOC). In other words, the Markov blanket is the smallest-dimension feature set 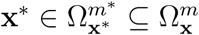, such that *P* (*y*|**x**) = *P* (*y*|**x**^∗^) [11].

For a BN, the Markov blanket contains only the parents, children, and other co-parents of children of the target node [11]. Markov blanket search algorithms allow for the identification of a Markov blanket set of a particular node (called a local network) without having to determine the entire BN (global network) [23]. Such algorithms perform a series of conditional independence tests on subsets of the data, accepting nodes into a Markov blanket set that are proven conditionally dependent on the target node given all other variables not in the Markov blanket set. PROMPTS utilizes the Grow-Shrink algorithm [24] to determine the Markov blanket of the DOC node (see materials and methods for details) and uses the Pomegranate [22] package in Python to perform BN structure search and training algorithms. Further computational details and links to all codes in the PROMPTS framework are provided in materials and methods.

Upon training to all pine litter data with the goal to predict DOC levels, the 32 features from the RFINN analysis were simplified to a BN model with only *m*^∗^ = 12 features. The resulting 12-feature BN model is depicted in Figure 2 and was found to be *Markov equivalent*^1^ to a Naive Bayes model. This specific BN model found through the structure search algorithm suggests that the abundance of each of the bacterial genera is conditionally independent of the rest, given the DOC target node [11]. Nodes for genera associated with high DOC levels are shaded blue and genera associated with low DOC levels are shown with green shaded nodes. The association of each genus with either high or low DOC levels was determined by conditioning on a particular genus and determining the marginal probability of DOC. It should be noted that the directions of arrows illustrate the independence assumptions made by the model and do not necessarily suggest that variation in DOC abundance drives variation in the abundance of the genera in the model.

**Fig 2.**
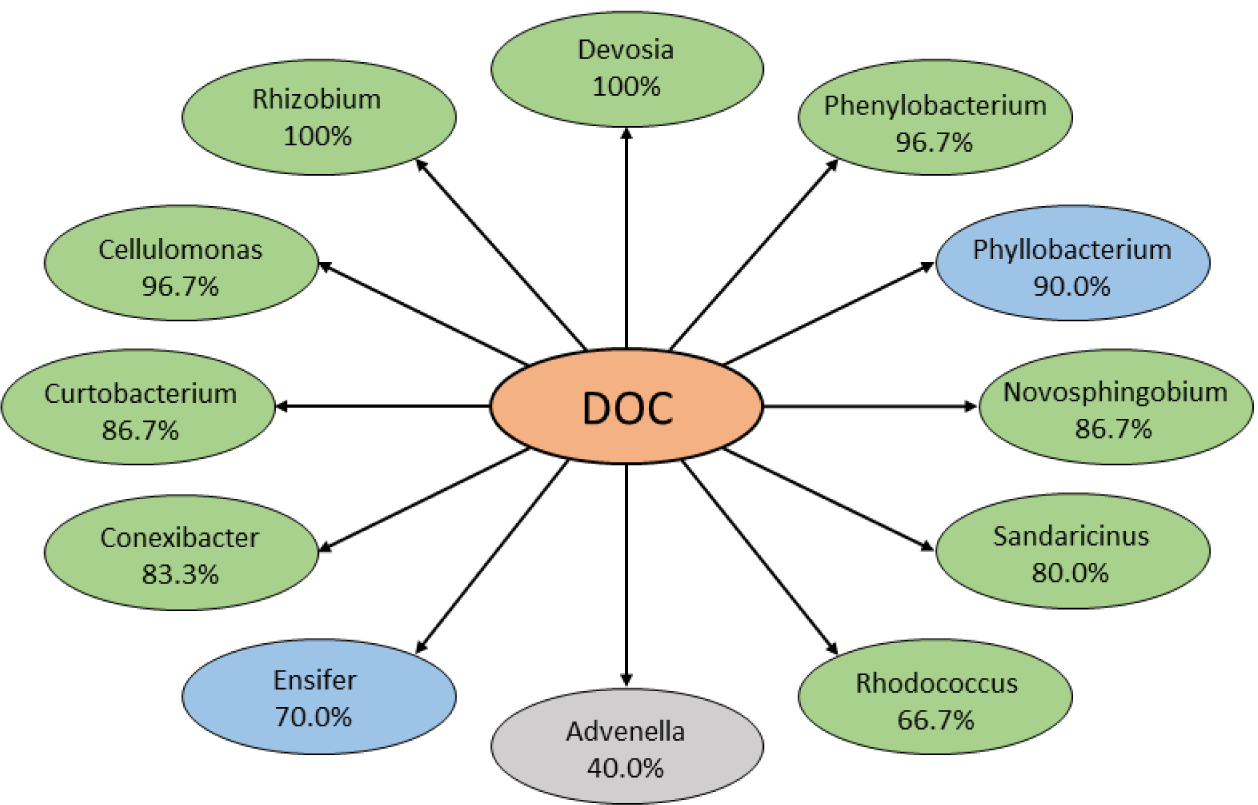
BN model of DOC using abundance of genera. The BN model composed of the Markov blanket of the DOC node is Markov equivalent to a Naive Bayes model in which all selected important taxa are conditionally independent given the DOC node. The Markov blanket set shown was determined using the entire set of pine samples. The central orange shaded node represents the target node, DOC, the blue shaded nodes represent genera associated with high DOC levels, and green shaded nodes represent genera associated with low DOC levels. The percent next to each genus indicates how frequently the genus was selected among 30 bootstrapped samples of ∼ 80% of the pine samples. Only taxa that appeared in at least 20 of the 30 Markov blanket sets were included in the final 11 feature model.

To probe the robustness of the Markov blanket identified in Figure 2 to variations within the training data sets, we next created 30 random data subsets, each containing ∼ 80% of the genera and DOC measurements from the full pine data set. For each random data subset, we repeated the Markov blanket identification procedure as outlined above. After identifying the Markov blanket sets of the 30 permutations, we ranked all features according to the frequency of their appearance within the Markov blanket sets (Figure 2). The frequency of appearance in bootstrap permutations for all Markov blanket features is shown in S1 Table. This bootstrapping analysis revealed that, of the 12 genera selected in the Markov blanket using the full data set, 11 genera were selected in at least 66.6% of the 30 Markov blanket sets determined from random data subsets. These 11 genera were also the most frequently selected features in the Markov blanket sets, and the frequency of these features ranged from 100% for the most frequent features (*Devosia, Rhizobium*) to 66.6% for the 11^th^ most frequent feature (*Rhodococcus*). DOC associations were consistent over all bootstrap permutations (e.g., genera associated with high DOC were always associated with high DOC in every permutation of the data).

### A simple Bayesian Network can accurately predict DOC from held-out 16S rRNA gene profile data collected from the same litter type

We next sought to validate whether a BN identified using PROMPTS could be expected to provide accurate quantitative predictions for testing data collected using the same litter type. Figure 1 illustrates the five-fold cross-validation approach that we used to perform this validation using the pine litter data. For each training permutation of the data, we repeated the BN identification tasks discussed above: First, we used RFINN for feature selection on each of the training data sets, and then we performed a Markov blanket search of the DOC node to construct a local BN that predicts high or low DOC. Next, we conducted a validation step in which we used the trained BN model to predict the DOC level for the held-out validation data. The results of this model validation (with five non-overlapping sets of 20% held-out data) are summarized in Figure 3A, which shows the receiving operating characteristic (ROC) curve representing prediction performance on held-out testing data for each permutation. For context, the area under the ROC curve (AUC) represents the probability of ranking a randomly selected high DOC sample higher than a randomly selected low DOC sample, where the model-generated rank is the predicted probability, p(DOC=High) [26]. The AUC ranged from 0.78 to 0.92, indicating the predictions were accurate for all permutations of the training and testing data. We next selected the permutation with the median AUC score of 0.86 (permutation 1, shown in dark blue in Figure 3A) as a representative train-test partition for subsequent in-depth analyses of the BN performance. Figure 3B presents the confusion matrix to visualize the predictive power of the BN on held-out testing data (similar confusion matrices for the other four permutations are shown in S1 FigureA-D). The upper left section of the confusion matrix shows the number of accurately identified low DOC samples, or *true negatives*, and the central block of the confusion matrix shows the number of accurately identified high DOC samples, or *true positives*. The lower right block shows the accuracy, where accuracy is defined as the number of true positives and true negatives divided by the total number of evaluated samples. Additional metrics such as the positive predictive value, which is the probability of a true positive, are shown in the remaining blocks.

**Fig 3.**
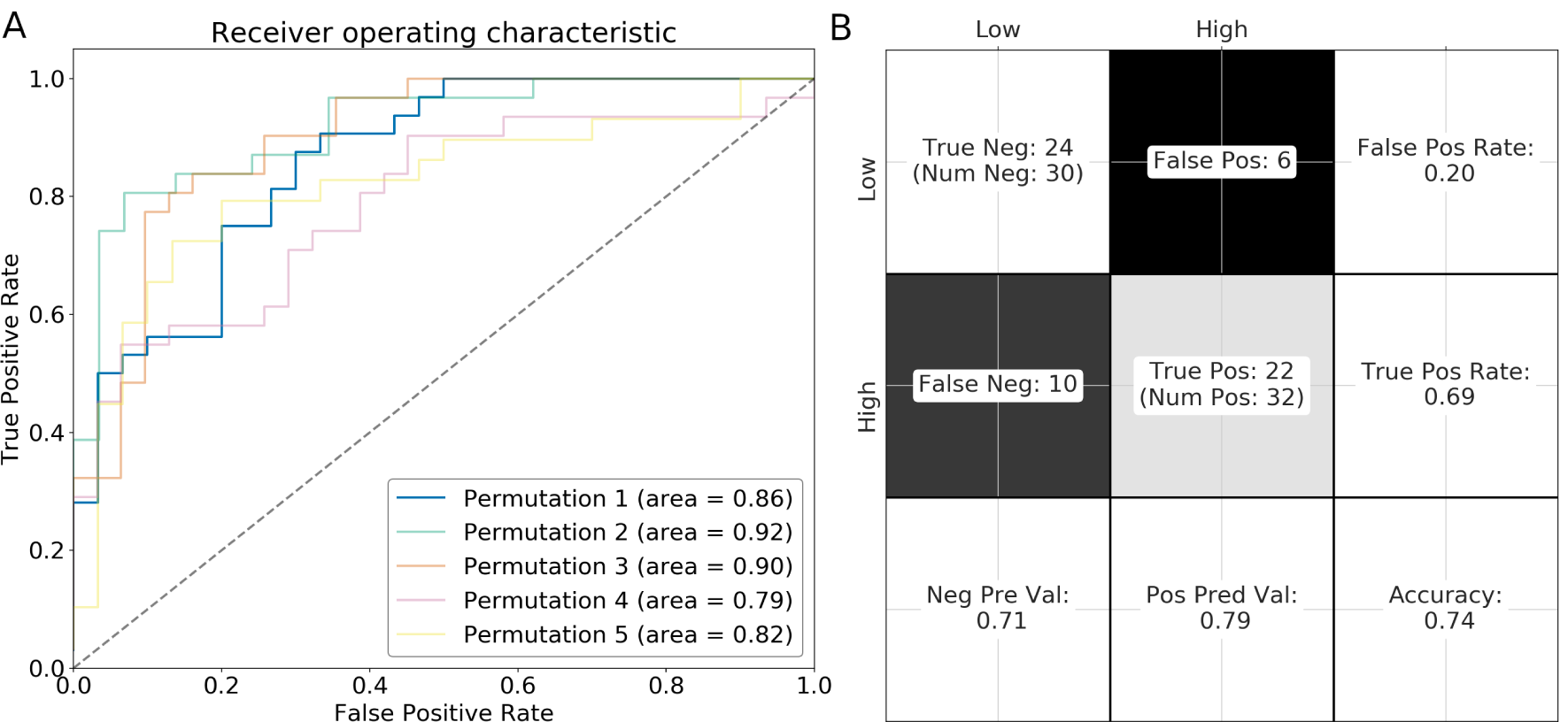
Prediction performance on held-out pine validation data. (A) The full data set was partitioned into five unique sets of training and testing data such that each set of held-out testing data was unique. The area under the ROC curve for each permutation ranged from 0.78 to 0.92 with a median of 0.86 corresponding to permutation 1. (B) A confusion matrix corresponding to prediction performance of the permutation 1 test data set shows 24 true negatives (upper left), and the central block shows 22 true positives. The lower right block shows that the permutation 1 prediction accuracy was 74%. See S1 Figure for similar confusion matrices for permutations 2-5.

For the confusion matrices shown in Figure 3B and S1 FigureA-D, every testing dataset that was predicted to have high-DOC with probability of 0.5 or more was classified as ‘High,’ and the remaining data sets were classified as ‘Low’. Although this simple binary interpretation demonstrates strong predictive results, it hides an important benefit of probabilistic predictions provided by the PROMPTS framework. To more carefully probe this benefit, we next asked how well the BN model could correctly distinguish between certain and uncertain DOC predictions. To illustrate this, Figure 4A shows a scatter plot of the actual DOC level versus the predicted probability of high DOC for held-out testing samples, i.e., *p*_*H*_ (**x**) := *p*(*DOC* = *High* | **x**). We defined an uncertainty cutoff of *η* to distinguish between certain and uncertain predictions, where predicted probability *p*_*H*_ (**x**) > *η* indicates a certain-High DOC prediction, and *p*_*H*_ (**x**) < 1 − *η* indicates a certain-Low DOC prediction. The cutoff *η* provides a simple mechanism to separate model predictions according to their expected accuracy: a small *η* prioritizes more predictions over uncertainty (e.g., takes all predictions at face value), whereas a large *η* weights the uncertainty of the model strongly (e.g., discards less trustworthy predictions).

**Fig 4.**
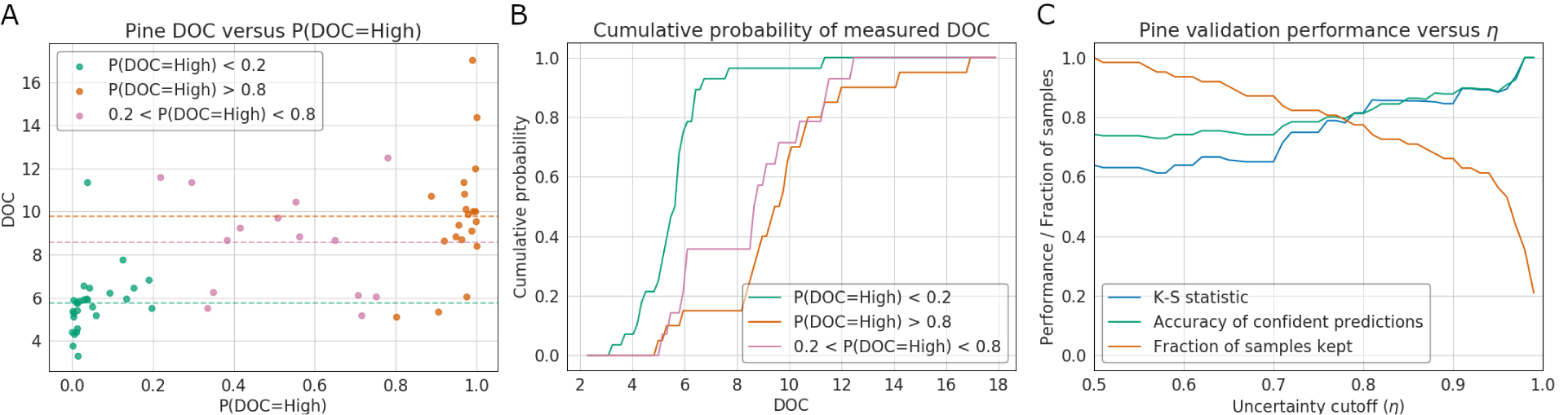
Classification and Uncertainty Quantification for DOC in Pine Litter. (A) Measured DOC versus predicted *P* (*DOC* = *High*) for held-out testing samples. Predictions with high certainty to have low DOC (P(DOC=High) < 0.2) and high DOC (P(DOC=High) > 0.8) are shown in green and orange, respectively. Low certainty predictions are shown in magenta. Dotted horizontal lines represent the average measured DOC for each group. (B) Cumulative distribution plot of the set of measured DOC values in the held-out testing set binned into certain-Low, uncertain, and certain-High DOC categories. The Kolmogorov-Smirnov (K-S) distance between true DOC values for certain-High and certain-Low samples was 0.814, p-value < 10^−3^. (C) K-S distance between certain-High and certain-Low samples (orange), fraction correct certain-High and certain-Low predictions (blue), and fraction of certain samples (cyan) versus uncertainty cut-off.

Figure 4A shows the probabilistic predictions of DOC levels for a cutoff of *η* = 0.8, where samples predicted with high certainty to have high and low DOC are labeled orange and green, respectively, and the remaining less certain predictions (i.e., where 1 − *η*≤ *p*_*H*_ (**x**) ≤ *η*) are shown in magenta. For the 62 sets of pine community testing data, 14 were classified as uncertain, and 48 communities were retained as certain. However, the communities that were classified as certain were strongly enriched for correct predictions (81%), as compared to the average accuracy for all communities (74%) or the accuracy for uncertain communities (50%), thus demonstrating that the BN is capable to correctly estimate which predictions are most likely to be accurate. The clear separation between certain-High and certain-Low DOC predictions is also shown in Figure 4B, which shows the cumulative probability distribution of measured DOC for samples classified as certain-Low (green), certain-High (orange) and uncertain (magenta). The two-sample Kolmogorov-Smirnov test was used to determine the statistical significance of the degree of separation in the distributions of measured DOC abundance corresponding to certain-Low and certain-High predictions. The cumulative probability distributions for the actual measured DOC levels had a Kolmogorov-Smirnov (K-S) distance of 0.814 between the certain-High and certain-Low classified samples (corresponding to p-value < 10^−7^), indicating that model’s distinctions between high and low DOC were strongly statistically significant. The effect of the probability cutoff *η* on model accuracy, K-S distance, and the fraction of certain predictions is shown in Figure 4C. The nearly monotonic increase in prediction accuracy with increasing *η* demonstrates that the model correctly assigns higher certainty to its more accurate predictions.

### Model predictions and uncertainty quantification prove accurate when transferred across variable litter types

Next, to investigate the transferability of the PROMPTS analyses, we tested the consensus 11 feature model (Figure 2) trained on the full pine data set to predict DOC outcomes of independent oak and grass litter data sets. The procedure for model training with pine data and validation with oak and grass data is summarized in Figure 1B. Prediction results from the same pipeline applied to a 2-feature model (with *Devosia* and *Rhizobium*) and the 12-feature model (with all Markov blanket features) are provided in supporting information (S2 Figure, S3 Figure, S4 Figure, S5 Figure). Figure 5A shows a scatter plot of measured DOC from the 99 held-out oak litter data samples versus the predicted probability of high DOC from the 11-feature BN trained on pine data. Using the same certainty cutoff as before (*η* = 0.8), DOC predictions were again categorized as certain-High (orange), certain-Low (green) and uncertain (magenta). Of the 99 total communities in the oak data, 35 predictions were discarded as uncertain, and 64 communities were classified as certain. Of those classified as certain, 83% were correctly predicted to be either below or above the median oak DOC, once again outperforming the average accuracy for all communities (74%) and the accuracy for uncertain communities (57%). As was the case for the pine litter validation data, Figure 5B again shows a clear separation between cumulative DOC distributions among the certain-High and certain-Low DOC communities with a K-S distance of 0.78 (p-value < 10^−8^) and Figure 5C shows that model accuracy improves as the certainty cut-off increases.

**Fig 5.**
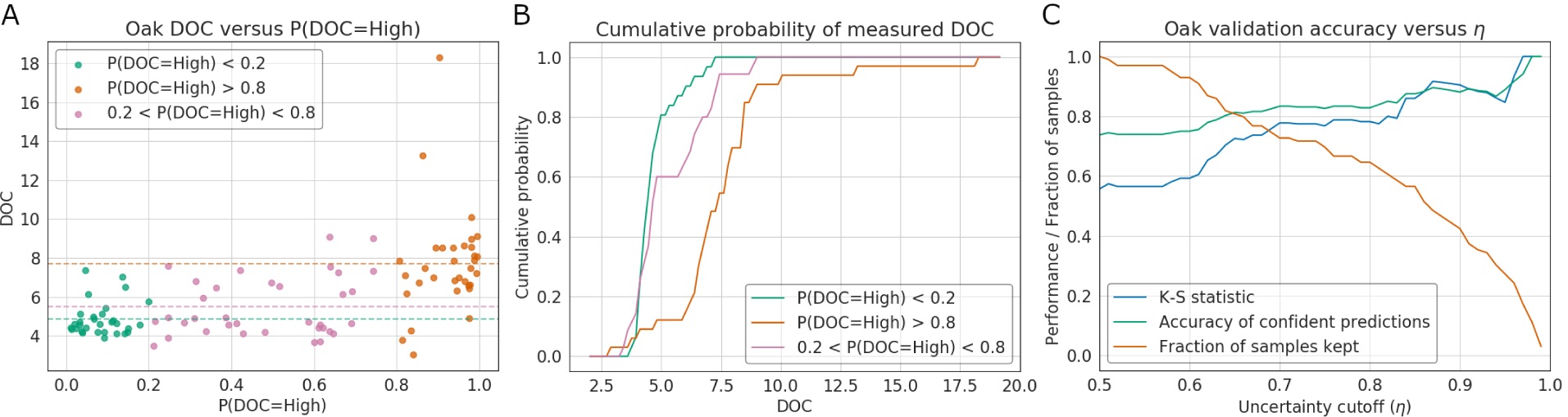
Cross-Litter Validation of DOC Classification in Oak Litter. (A) Oak DOC versus predicted *P* (*DOC* = *High*). Samples predicted with high confidence to have low DOC (P(DOC=High) < 0.2) and high DOC (P(DOC=High) > 0.8) are shown in green and orange, respectively. Predictions for which confidence is less certain are shown in magenta. Dotted horizontal lines represent the average measured DOC for each group.(B) The set of measured DOC values in the oak data set binned into low, uncertain, and high DOC categories according to the predicted probability of high DOC using a BN trained on pine samples. The Kolmogorov-Smirnov (K-S) distance between true DOC values of samples with a high predicted DOC and DOC values of samples with low predicted DOC was 0.78 (p-value < 10^−8^). (C) K-S distance between certain-High and certain-Low samples (orange), fraction correct certain-High and certain-Low predictions (blue), and fraction of certain samples (cyan) versus uncertainty cut-off.

To further investigate the transferability of the BN model, we applied the same model trained on pine data to samples from a grass litter decomposition experiment (Figure 6A,B,C). Although the model was able to predict DOC for grass litter samples with statistically significant accuracy (K-S distance = 0.58, p value = 0.030), prediction performance was notably worse than with the oak litter data set. Of the 99 total communities in the grass data, 34 predictions were discarded as uncertain, and 65 communities were classified as certain. Of those classified as certain, 60% were correctly predicted to be either below or above the median grass DOC, once again outperforming the average accuracy for all communities (54%) and the accuracy for uncertain communities (41%). Poor performance on the grass data set represents an important failure case, demonstrating that the model loses relevance when applied to disparate litter types. However, we note that in all testing circumstances, prediction accuracy improves as the certainty cut-off increases, suggesting that the analysis is correctly able to identify which of its predictions are more likely to be accurate.

**Fig 6.**
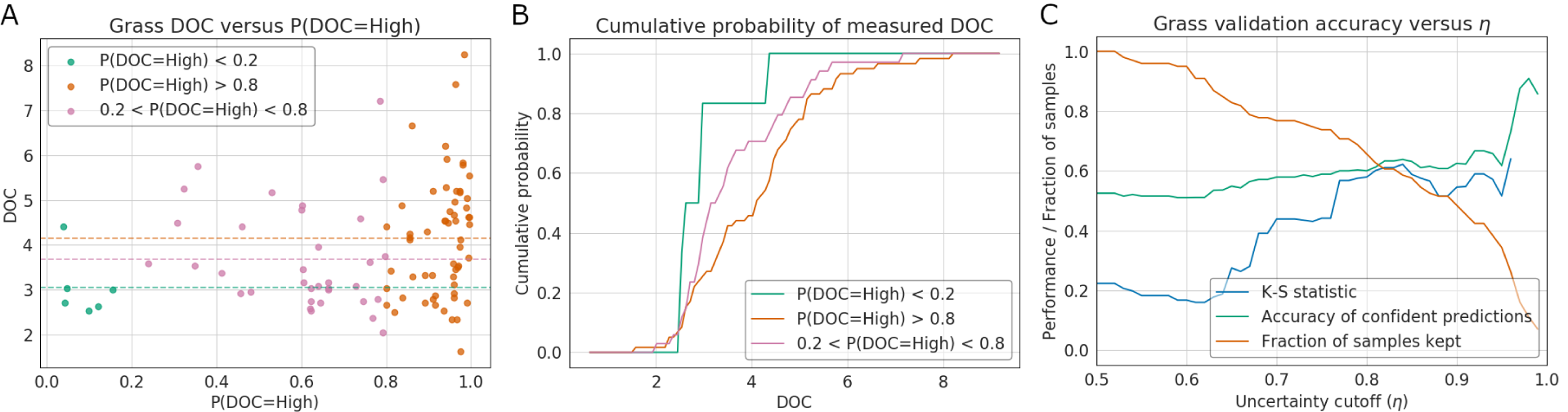
Grass DOC versus predicted *P* (*DOC* = *High*). (A) The average DOC of samples predicted to have low DOC (P(DOC=High) < 0.2), high DOC (P(DOC=High) > 0.8), and uncertain samples are shown as green, orange, and magenta dotted horizontal lines.(B) The set of measured DOC values in the grass data set binned into low, uncertain, and high DOC categories according to the predicted probability of high DOC using a BN trained on pine samples. The Kolmogorov-Smirnov (K-S) distance between true DOC values of samples with a high predicted DOC and DOC values of samples with low predicted DOC was 0.58, p value = 0.030. (C) K-S distance between certain-High and certain-Low samples (orange), fraction correct certain-High and certain-Low predictions (blue), and fraction of certain samples (cyan) versus uncertainty cut-off.

Finally, we sought to explore both in-study and cross-study prediction accuracy on held-out data from pine, oak, and grass data sets using a series of BN models trained on combinations of pine, oak, and grass litter training data. This validation analysis included seven combinations of training scenarios: only pine data, only oak data, only grass data, pine and oak data, pine and grass data, oak and grass data, and pine, oak, and grass data. Using genera that appeared in the intersection set in pine, oak, and grass data sets, model training and prediction was performed over 50 bootstrap permutations in which ∼ 20% of each data set was randomly selected and withheld for testing. Prediction performance was assessed using the K-S distance between measured DOC samples categorized into certain-High and certain-Low sample sets based on the average “out of bag” model predictions using an uncertainty cutoff *η* = 0.8. This comprehensive validation analysis revealed significant prediction performance in all training and testing circumstances, and that a BN trained on pine, oak, and grass data serves as a general model that accurately estimates DOC on held-out samples from each litter type (Figure 7). Cumulative distribution plots of measured DOC separated according to out of bag DOC predictions for every training and testing scenario are shown in S6 Figure. Furthermore, this analysis showed that oak data is always well predicted and that model predictions are generally improved when the oak data set is included in the training data. The generality of the oak data set is supported by examining the selected features from each data set, where several of the most frequently selected taxa in pine (e.g., *Rhizobium* and *Ensifer*) and grass (e.g., *Shinella* and *Stenotrophomonas*) were also selected in the oak data set, but were not selected among all three data sets. The frequency with which features appeared in each Markov blanket set and their associations with DOC over the 50 bootstrap permutations after training on pine, oak, and grass data is shown in S2 Table.

**Fig 7.**
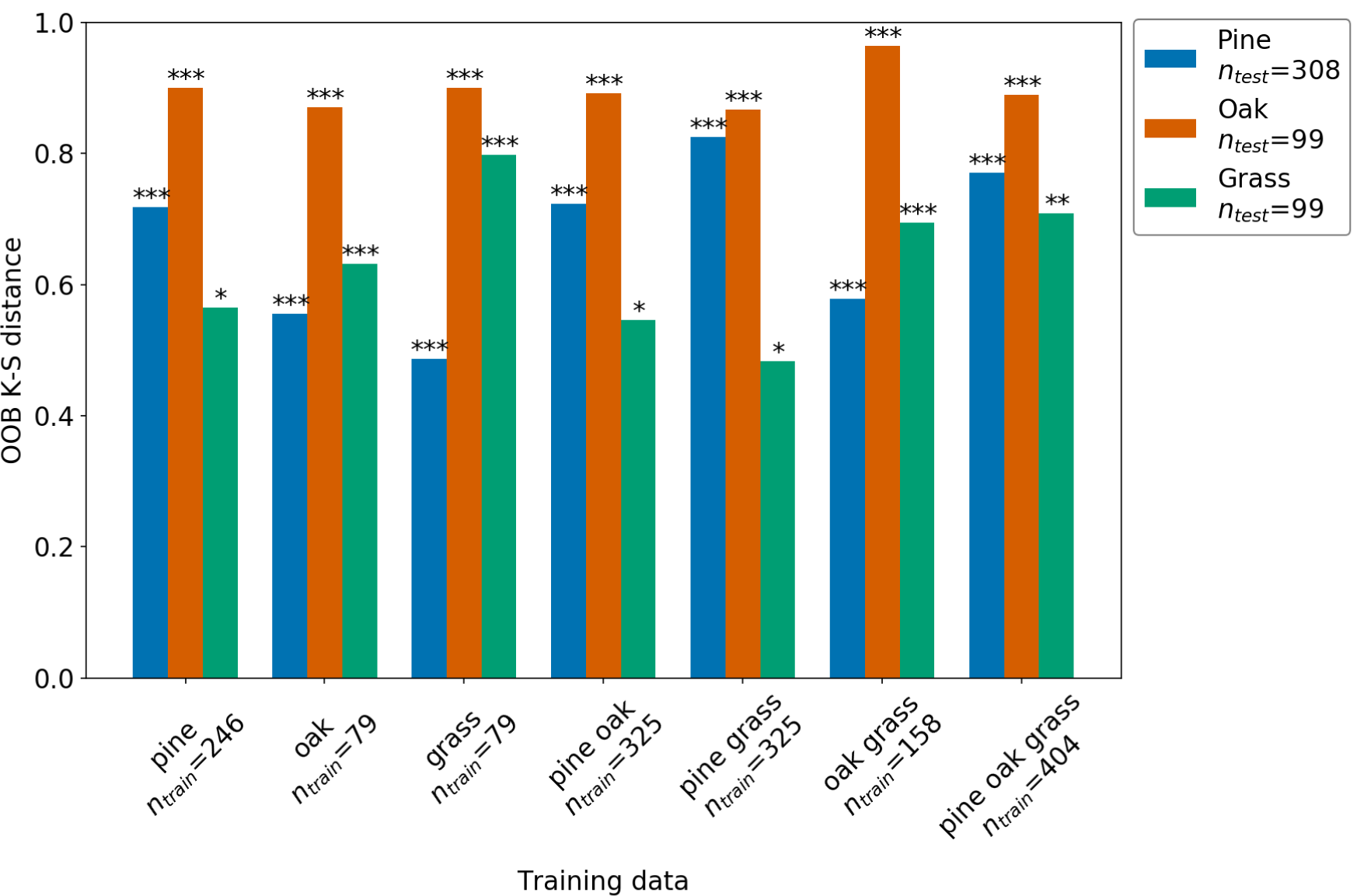
Out of bag prediction performance on pine, oak, and grass data sets using models trained on combinations of pine, oak, and grass data. The pine, oak, and grass data sets were partitioned into training and testing data over 50 boot-strap permutations with each permutation reserving 20% of samples for held-out testing. For each permutation, a set of Bayesian network models were trained using seven different data groups: only pine, only oak, only grass, pine and oak, pine and grass, oak and grass, and pine, oak, and grass data. Using the average prediction (“out of bag” or “oob” prediction) for each pine (blue), oak (orange) and grass (green) sample, the K-S test was used to assess model prediction performance and significance. Sample size, *n*_*train*_ and *n*_*test*_, is indicated for each training or testing group, respectively. The bars show the K-S statistics (vertical height) and associated p-value: ^∗^ < 0.05, ^∗∗^ < 0.01, and ^∗∗∗^ < 0.001.

## Discussion

By coupling the rigorous selection of biologically interpretable features with the ability to make quantitative and probabilistic predictions, Bayesian networks (BNs) offer a powerful tool for the computational analysis of complex microbial communities. In this study, we created the analysis package PROMPTS to constrain BNs and infer a probabilistic graphical model that captures the interactions between relative microbial abundance and dissolved organic carbon (DOC) in a series of pine litter decomposition experiments, each starting from a different inoculum composition. By determining the Markov blanket of the DOC node in the resulting BN, PROMPTS identified a small set of specific bacterial genera whose relative abundances directly constrain the probability of high or low DOC levels. In addition to using cross-validation to verify that the BN could predict DOC within held-out pine litter decomposition experiments, we also verified that the PROMPTS analysis trained on pine data could predict DOC outcomes for independent decomposition experiments in oak and grass litter. To our knowledge, this study presents the first example of a validated probabilistic graphical model approach that predicts microbiome function across multiple substrate environments.

The first major benefit of the PROMPTS framework analyses is the ability to identify a small number of important and biologically interpretable features that robustly correlate with collective community behavior. By determining the Markov blanket of the DOC node using many permutations of training data, our BN analysis rigorously searched over all features to select a minimal set of microbial genera that constrain the probability of high DOC abundance. Several of the genera identified in the Markov blanket set (Figure 2) are known to be involved in litter decomposition either directly or indirectly [2, 27–33]. Of the 11 most robustly selected features, *Phyllobacterium* (selected in 90% of BNs) and *Ensifer* (selected in 70% of BNs) were the only two genera associated with high DOC; these known nitrogen-fixing bacteria are likely helpful to promote decomposition of pine litter, which is often nitrogen limited [2, 29, 33]. Of the remaining genera associated with low DOC, *Devosia* (100% of BNs), *Cellulomonas* (97% of BNs), and *Novosphingobium* (87% of BNs), are all known to degrade cellulose, hemicellulose, and/or lignin, which make up the main components of plant litter [2, 27, 34, 35]. These genera are likely playing a direct role in plant litter decomposition, while the *Rhizobium* genus (100% of BNs) is indirectly involved by providing nitrogen and phosphorus to lignocellulose degrading microorganisms [31]. Using the key genera identified in the analysis of pine litter decomposition data, PROMPTS could accurately classify high and low DOC samples for oak litter (p-value < 10^−8^) and grass litter (p-value =0.030) decomposition experiments. These strong predictions suggest that relationships between the abundances of key bacterial genera and DOC are robust, even across litter types. Moreover, this result of transference in composition-to-function predictability from pine to oak litter supports the hypothesis that common microbial drivers of DOC abundance can occur in diverse ecosystems despite the strong role that litter chemistry has in shaping distinctive decomposer microbiomes [2]. Identifying small sets of genera whose abundances are robustly predictive of collective community functions (e.g., DOC level) provides a necessary first step towards designing model microbial communities to control those functions (e.g., to promote increased organic carbon accumulation), even when faced with uncertain environments (e.g., different litter substrates).

The second major benefit of the PROMPTS analysis framework is the ability to evaluate predictions of community-level behaviors with the rigorous quantification of expected prediction uncertainties. In all testing circumstances, the BN model predicted DOC outcomes of confidently predicted samples with greater accuracy compared to uncertain predictions. This trend is clearly demonstrated in Figures 4C, 5C, and 6C, which all show that as the prediction uncertainty cut-off, *η*, is increased from *η* = 0.5 (no uncertainty selection) to *η* = 0.95 (strong selection), the accuracy of predictions increases for all three litter types. Consequently, this cross-validation on independent data sets not only demonstrates that the PROMPTS model can successfully make accurate predictions when applied beyond its training circumstances, but that the model also correctly assigns greater collective uncertainty to its less accurate predictions. Such models, once validated to make accurate estimates of their own uncertainty, could be especially useful not only for engineering design (e.g., communities with the most certain high DOC predictions are most likely to succeed to increase carbon levels in soil), but also for subsequent engineering research (e.g., careful testing and analysis of uncertain predictions may help to discover secondary interactions that are not yet confidently characterized in the training data).

The approach to train and cross-validate BNs that characterize the interplay of microbial abundances and quantified community-level behavior provides a robust and interpretable taxonomic ‘parts list’ that constrains probabilistic predictions of collective community behaviors. These insights from structured predictive models help set the stage for future applications to understand and reengineer microbial communities toward specific industrial or biomedical goals. To enable such applications, we envision future efforts that seek to augment the predictive ability of PROMPTS with more complex mechanistic understanding. Indeed, probabilistic graphical model (PGM) based approaches such as SPIEC-EASI [10] and MERLIN [9] have recently been developed that offer a promising approach for comprehensive reconstruction of interaction networks. Designed to address the sparse and compositional nature of microbial abundance data, SPIEC-EASI has shown promise for inferring microbial interaction networks compared to frequently used [36] correlation-based techniques. MERLIN has proven more effective than BNs for determining interaction networks on synthetic gene expression data but has not yet been applied to model microbial communities. Unlike the BN approach presented in this study, these methods have not been bench-marked for their ability to make accurate predictions of target variables, but could be used in addition to models validated for prediction accuracy. Ultimately, the combination of validated prediction models with reconstructed interaction networks could provide a powerful engineering tool to guide the strategic optimization of microbiomes.

## Materials and methods

### Plant litter decomposition experiments

Plant litter decomposition experiments were performed in laboratory microcosms consisting of 125ml serum bottles containing 7g of sand and 0.12g of plant leaf litter (either Ponderosa pine needles, scrub oak leaves, or a 50:50 mix of two grasses (Stipa and Hilaria) common in the U.S. southwest). The microcosms were inoculated with soil microbial communities, sealed with crimp caps, and incubated at 25°C for about 6 wks. Cumulative CO_2_ was determined from repeated measurements over the incubation period and the final abundance of DOC was measured at the end of the incubation. For the pine experiment, “low” and “high” DOC communities were delineated as the left and right thirds (approximately) of the observed DOC distribution, and DNA representing these microcosm community cohorts was extracted for community analysis. The communities were taxonomically profiled by PCR amplification and Illumina sequencing of fungal and bacteria ribosomal RNA gene fragments. The pine litter experiment was described in detail [5]. The oak and grass litter experiments were performed similar to the pine litter experiment, with the following modifications: only 100 (not 206) source soil communities were used to inoculate microcosms, and each source soil community was inoculated into 2 replicate (not 3) microcosms.

### Data pre-processing

The pine data were rarefied such that OTU abundance in each sample totaled to 1026 counts, and the oak and grass genera were rarefied such that OTU abundance in each sample summed to 1023 counts. The oak and grass OTU tables were normalized to match the pine data composition so that OTU abundance of each sample summed to 1026 counts. For all analyses, taxa were evaluated at the genus taxonomic rank. To calculate relative genera abundance, OTUs classified with greater than 70% confidence for a particular genus were summed together. For validation on held-out data, the pine data set of 308 samples was divided according to a K-fold partitioning scheme with 5-fold permutations of training and testing data. K-fold partitioning of the data set ensured that each set of testing samples was unique so that all samples were subject to held-out testing. In each permutation, replicate samples were kept within either training or testing data sets to ensure independence between sets. Genera abundance was binned into categorical variables representing low, medium, and high abundance levels based on training data statistics (see next section for detail). Similarly, DOC was binned into high or low categorical variables based on the median DOC in the training data. Genera abundance in the oak and grass datasets were binned into categorical variables based on the abundance of taxa in the pine litter (training) dataset. Measured oak and grass DOC was not binned, but left as a continuous variable that was compared to the predicted probability of high DOC.

### Markov blanket search

We applied the Grow-Shrink algorithm [24] to determine the Markov blanket of the DOC node. Defining 𝒰 as the set of genera abundance and DOC, 𝒰 = {**x**, *y*}, the Grow-shrink algorithm to determine the Markov blanket of *Y* ∈ 𝒰 is shown in Procedure 1.

#### Procedure 1: Grow-Shrink algorithm for determining the Markov blanket

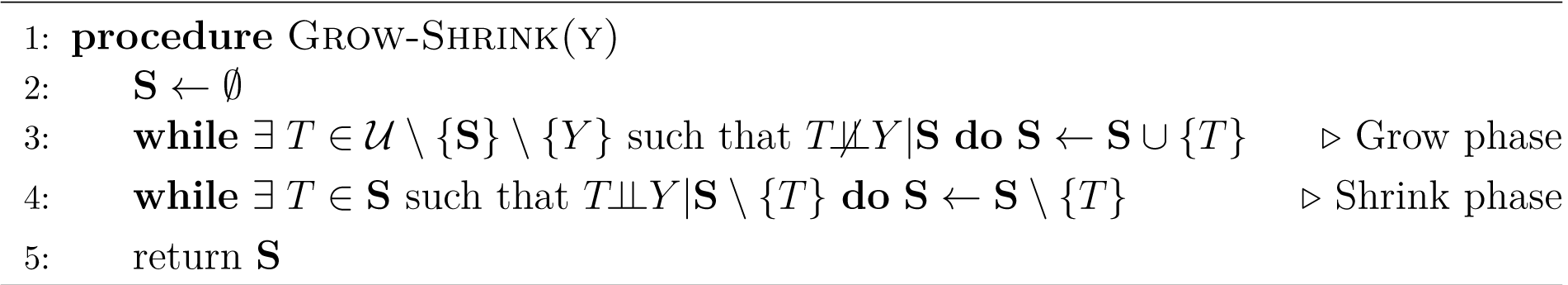

To perform conditional dependence tests, variables are tested for d-separation. Given a set of variables **S** ∈ 𝒰, two variables *T* ∈ 𝒰 and *Y*∈ 𝒰 are d-separated if every path connecting variables is blocked according to the following criteria: If arrows on the path connecting variables *T* and *Y* meet either head to tail or tail to tail at a node in the set **S**, or if the arrows on the path meet head to head at a node not in **S** and none of the node’s descendants are in **S**. Variables *T* and *Y* are conditionally independent given the set **S** if the variables *T* and *Y* are d-separated given the set **S** [7, 23].

### Evaluating prediction performance

The Kolmogorov-Smirnov (K-S) two sample test was performed to provide an easily interpreted metric of prediction performance. The null hypothesis of the two sample K-S test is that the cumulative distributions of observations in each sample are equivalent. This statistic was applied to determine if high and low DOC predictions separated measured DOC into distinct groups; if the p value is statistically significant, then the Bayesian network makes predictions of DOC levels that are collectively statistically significant. The test statistic is defined as

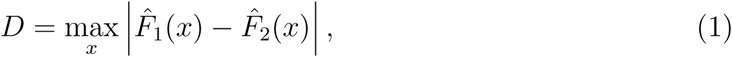

where 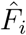 is the empirical cumulative distribution function of the *i*^*th*^ sample. The test statistic, *D*, ranges from 0 to 1 and represents the maximum distance between two cumulative distributions. The p value of the K-S test represents the probability of observing the test statistic, *D*, if the two samples were collected from identical distributions [37]. The two sample K-S test statistic and p value were calculated using the default settings of Scipy’s ks_2samp function [38].

## Data and Code Availability

All data and codes are available at https://github.com/MunskyGroup/PROMPTS.

## Acknowledgments

JT, RJ, JD and BM were supported by grant F255LANL2018 from the U.S. Department of Energy Office of Biological and Environmental Research, Genomic Science program. BM was also supported by National Institutes of Health (R35 GM124747). NL was supported by the U.S. Department of Energy through the Laboratory Directed Research and Development (LDRD) program at Los Alamos National Laboratory. The funders had no role in study design, data collection and analysis, decision to publish, or preparation of the manuscript.

## Author contributions

Conceptualization, BM, JD, JT and NL; Methodology, JT, BM and NL; Experiments: JF, RJ, RD, and MEK; Writing Original Draft: JT, NL, and BM; Writing Review & Editing, JT, NL, JD, BM, and MEK; Funding Acquisition: JD and BM; Resources: JD, RJ and RD; Supervision: JD and BM.

## Supporting information

**S1 Table. Markov blanket feature selection table for pine.** A Markov blanket search of the DOC node using the 32 features selected by RFINN was performed over 30 bootstrap iterations of 80% of the pine data set. The blue genera were found to be in the Markov blanket set of the DOC node after performing a Markov blanket search using the entire pine data set. The prevalence column shows how frequently the genera were selected in the 30 bootstrap iterations, where Devosia and Rhizobium were selected in every iteration. DOC associations for each feature are determined by conditioning the Bayesian network on the feature and marginalizing over the rest. Only the top 20 most prevalent features are shown.

**S2 Table. Markov blanket feature selection table for pine, oak, and grass** The PROMPTS framework was applied over 50 bootstrap permutations of training data pulled from pine, oak, and grass litter datasets. The frequency of appearance of each feature after training on pine, oak, and grass data is shown in each respective column. The DOC Assoc. columns shows whether each genera was associated with High or Low DOC according to the parameters of the trained BN model.

**S1 Figure. Confusion matrices of test permutations 2-5.** Entries in row *i* and column *j* of each confusion matrix represent the number of samples in class *i* predicted by the model to be in class *j*. As shown here, the upper left section of the confusion matrix shows the number of accurately identified low DOC samples, or *true negatives*, and the central block of the confusion matrix shows the number of accurately identified high DOC samples, or *true positives*. The lower right block shows the accuracy, where accuracy is defined as the number of true positives and true negatives divided by the total number of evaluated samples. Additional metrics such as the positive predictive value, which is the probability of a true positive, are shown in the remaining blocks.

**S2 Figure. Prediction performance of two-feature model on oak data** (A) Samples predicted with high confidence to have low DOC (P(DOC=High) < 0.2) and high DOC (P(DOC=High) > 0.8) are shown in green and orange, respectively. Predictions for which confidence is less certain are shown in magenta. Dotted horizontal lines represent the average measured DOC for each group.(B) The set of measured DOC values in the oak data set binned into low, uncertain, and high DOC categories according to the predicted probability of high DOC using a BN trained on pine samples.

**S3 Figure. Prediction performance of 12-feature model on oak data** (A) Samples predicted with high confidence to have low DOC (P(DOC=High) < 0.2) and high DOC (P(DOC=High) > 0.8) are shown in green and orange, respectively. Predictions for which confidence is less certain are shown in magenta. Dotted horizontal lines represent the average measured DOC for each group. (B) The set of measured DOC values in the oak data set binned into low, uncertain, and high DOC categories according to the predicted probability of high DOC using a BN trained on pine samples.

**S4 Figure. Prediction performance of two-feature model on grass data** (A) Samples predicted with high confidence to have low DOC (P(DOC=High) < 0.2) and high DOC (P(DOC=High) > 0.8) are shown in green and orange, respectively. Predictions for which confidence is less certain are shown in magenta. Dotted horizontal lines represent the average measured DOC for each group. (B) The set of measured DOC values in the oak data set binned into low, uncertain, and high DOC categories according to the predicted probability of high DOC using a BN trained on pine samples.

**S5 Figure. Prediction performance of 12-feature model on grass data** (A) Samples predicted with high confidence to have low DOC (P(DOC=High) < 0.2) and high DOC (P(DOC=High) > 0.8) are shown in green and orange, respectively. Predictions for which confidence is less certain are shown in magenta. Dotted horizontal lines represent the average measured DOC for each group. (B) The set of measured DOC values in the oak data set binned into low, uncertain, and high DOC categories according to the predicted probability of high DOC using a BN trained on pine samples.

**S6 Figure. Cumulative distribution plots of out of bag predictions on pine, oak, and grass data sets using models trained on combinations of pine, oak, and grass data.** Using genera that appeared in the intersection set in pine, oak, and grass data sets, model training and prediction was performed over 50 bootstrap permutations in which ∼ 20% of each data set was randomly selected and withheld for testing. Each row indicates the training data set, and each column corresponds to the testing data set.

## Footnotes

1 Two structures with different edge directions are said to be *Markov equivalent* if both satisfy equivalent independence assumptions [25].

