## Supplemental Tables and Figures for "Probabilistic Ranking Of Microbiomes Plus Taxa Selection to discover and validate microbiome function models for multiple litter decomposition studies"

### S1 Table. Markov Blanket Feature Selection Table

| Top Features | Prevalence | DOC association |
|---|---|---|
| Devosia | 30 | Low |
| Rhizobium | 30 | Low |
| Phenylobacterium | 29 | Low |
| Cellulomonas | 29 | Low |
| Phyllobacterium | 27 | High |
| Curtobacterium | 26 | Low |
| Novosphingobium | 26 | Low |
| Conexibacter | 25 | Low |
| Sandaracinus | 24 | Low |
| Ensifer | 21 | High |
| Rhodococcus | 20 | Low |
| Modestobacter | 16 | Low |
| Aureimonas | 13 | High |
| Advenella | 12 | Low |
| Bradyrhizobium | 8 | Low |
| Sphingomonas | 7 | High |
| Sphingobium | 5 | Low |
| Oxalicibacterium | 4 | High |
| Achromobacter | 3 | Low |
| Opitutus | 3 | Low |

A Markov blanket search of the DOC node using the 32 features selected by RFINN was performed over 30 bootstrap iterations of 80% of the pine data set. The blue genera were found to be in the Markov blanket set of the DOC node after performing a Markov blanket search using the entire pine data set. The prevalence column shows how frequently the genera were selected in the 30 bootstrap iterations, where Devosia and Rhizobium were selected in every iteration. DOC associations for each feature are determined by conditioning the Bayesian network on the feature and marginalizing over the rest. Only the top 20 most prevalent features are shown.

#### S2 Table. Markov blanket feature selection table for pine, oak, and grass

| | Pine | | Oak | | Grass | |
|---|---|---|---|---|---|---|
| Genera | MB Counts | DOC Assoc. | MB Counts | DOC Assoc. | MB Counts | DOC Assoc. |
| Devosia | 50.0 | Low | 10.0 | Low | 11.0 | Low |
| Rhizobium | 49.0 | Low | 50.0 | Low | – | – |
| Cellulomonas | 45.0 | Low | 2.0 | Low | – | – |
| Ensifer | 42.0 | High | 26.0 | High | – | – |
| Rhodococcus | 39.0 | Low | – | – | – | – |
| Curtobacterium | 33.0 | Low | 3.0 | High | – | – |
| Phyllobacterium | 31.0 | High | – | – | – | – |
| Flavobacterium | 29.0 | Low | – | – | 9.0 | Low |
| Sphingomonas | 28.0 | High | 1.0 | High | – | – |
| Conexibacter | 26.0 | Low | – | – | – | – |
| Phenylobacterium | 25.0 | Low | – | – | – | – |
| Altererythrobacter | 22.0 | High | – | – | – | – |
| Novosphingobium | 19.0 | Low | 5.0 | Low | – | – |
| Modestobacter | 18.0 | Low | 13.0 | High | – | – |
| Georgenia | 16.0 | High | – | – | – | – |
| Pedomicrobium | 11.0 | High | – | – | – | – |
| Agaricicola | 11.0 | Low | – | – | – | – |
| Chitinophaga | 10.0 | Low | – | – | – | – |
| Aureimonas | 10.0 | High | – | – | – | – |
| WPS-1 genera incertae sedis | 7.0 | Low | 19.0 | Low | – | – |
| Bradyrhizobium | 6.0 | Low | 48.0 | Low | – | – |
| Sphingobium | 5.0 | Low | – | – | – | – |
| Achromobacter | 5.0 | Low | 7.0 | Low | – | – |
| Pigmentiphaga | 4.0 | Low | – | – | – | – |
| Azospirillum | 4.0 | High | – | – | – | – |
| Pedobacter | 4.0 | Low | – | – | – | – |
| Opitutus | 4.0 | Low | – | – | 3.0 | Low |
| Leifsonia | 3.0 | High | – | – | – | – |
| Sandarakinorhabdus | 3.0 | High | – | – | – | – |
| Roseomonas | 3.0 | High | – | – | – | – |
| Halomonas | 2.0 | High | – | – | – | – |
| Aquabacterium | 2.0 | High | – | – | – | – |
| Blastococcus | 2.0 | Low | 23.0 | High | – | – |
| Belnapia | 1.0 | High | – | – | – | – |
| Neorhizobium | 1.0 | High | – | – | – | – |
| Aminobacter | 1.0 | Low | – | – | 2.0 | Low |
| Nocardioides | 1.0 | Low | 1.0 | High | – | – |
| Sandaracinus | 1.0 | Low | – | – | – | – |
| Spartobacteria genera incertae sedis | 1.0 | Low | 35.0 | Low | 10.0 | Low |
| Steroidobacter | 1.0 | Low | – | – | – | – |
| Glaciimonas | 1.0 | High | 1.0 | High | – | – |
| Microvirga | – | – | 34.0 | High | – | – |
| Luteolibacter | – | – | 20.0 | Low | – | – |
| Arthrobacter | – | – | 9.0 | High | – | – |
| Hydrogenophaga | – | – | 8.0 | Low | 3.0 | Low |
| Pontibacter | – | – | 5.0 | High | – | – |
| Promicromonospora | – | – | 5.0 | High | – | – |
| Pseudomonas | – | – | 5.0 | Low | – | – |
| Domibacillus | – | – | 4.0 | High | – | – |
| Shinella | – | – | 4.0 | Low | 47.0 | Low |
| Stenotrophomonas | – | – | 2.0 | Low | 44.0 | Low |
| Agrococcus | – | – | 2.0 | High | – | – |
| Caulobacter | – | – | 1.0 | Low | – | – |
| Microbacterium | – | – | 1.0 | High | – | – |
| Methylobacillus | – | – | 1.0 | High | – | – |
| Adhaeribacter | – | – | 1.0 | High | – | – |
| Luteibacter | – | – | 1.0 | Low | – | – |
| Algoriphagus | – | – | 1.0 | High | – | – |
| Mycetocola | – | – | 1.0 | High | – | – |
| Brevundimonas | – | – | – | – | 13.0 | Low |
| Massilia | – | – | – | – | 3.0 | Low |

The PROMPTS framework was applied over 50 bootstrap permutations of training data pulled from pine, oak, and grass litter datasets. The frequency of appearance of each feature after training on pine, oak, and grass data is shown in each respective column. The DOC Assoc. columns shows whether each genera was associated with High or Low DOC according to the parameters of the trained BN model.

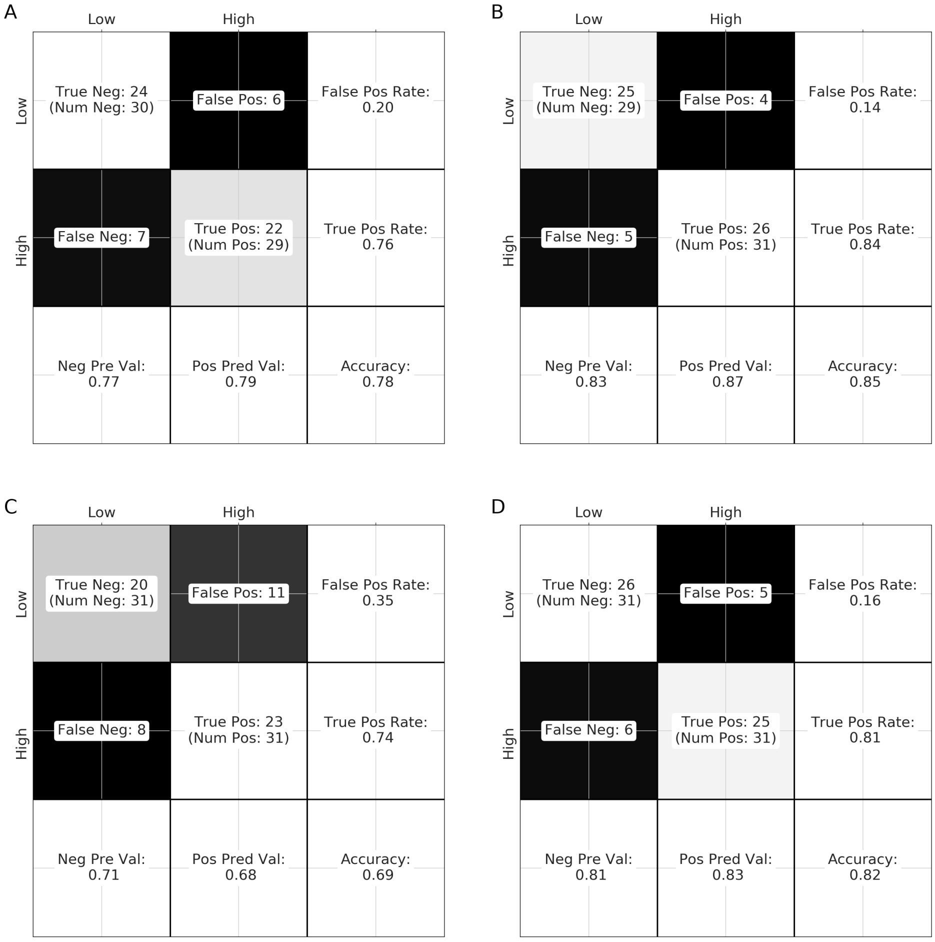


**S1 Figure. Confusion matrices of test permutations 2-5.** Entries in row $i$ and column $j$ of each confusion matrix represent the number of samples in class $i$ predicted by the model to be in class $j$. As shown here, the upper left section of the confusion matrix shows the number of accurately identified low DOC samples, or *true negatives*, and the central block of the confusion matrix shows the number of accurately identified high DOC samples, or *true positives.* The lower right block shows the accuracy, where accuracy is defined as the number of true positives and true negatives divided by the total number of evaluated samples. Additional metrics such as the positive predictive value, which is the probability of a true positive, are shown in the remaining blocks.

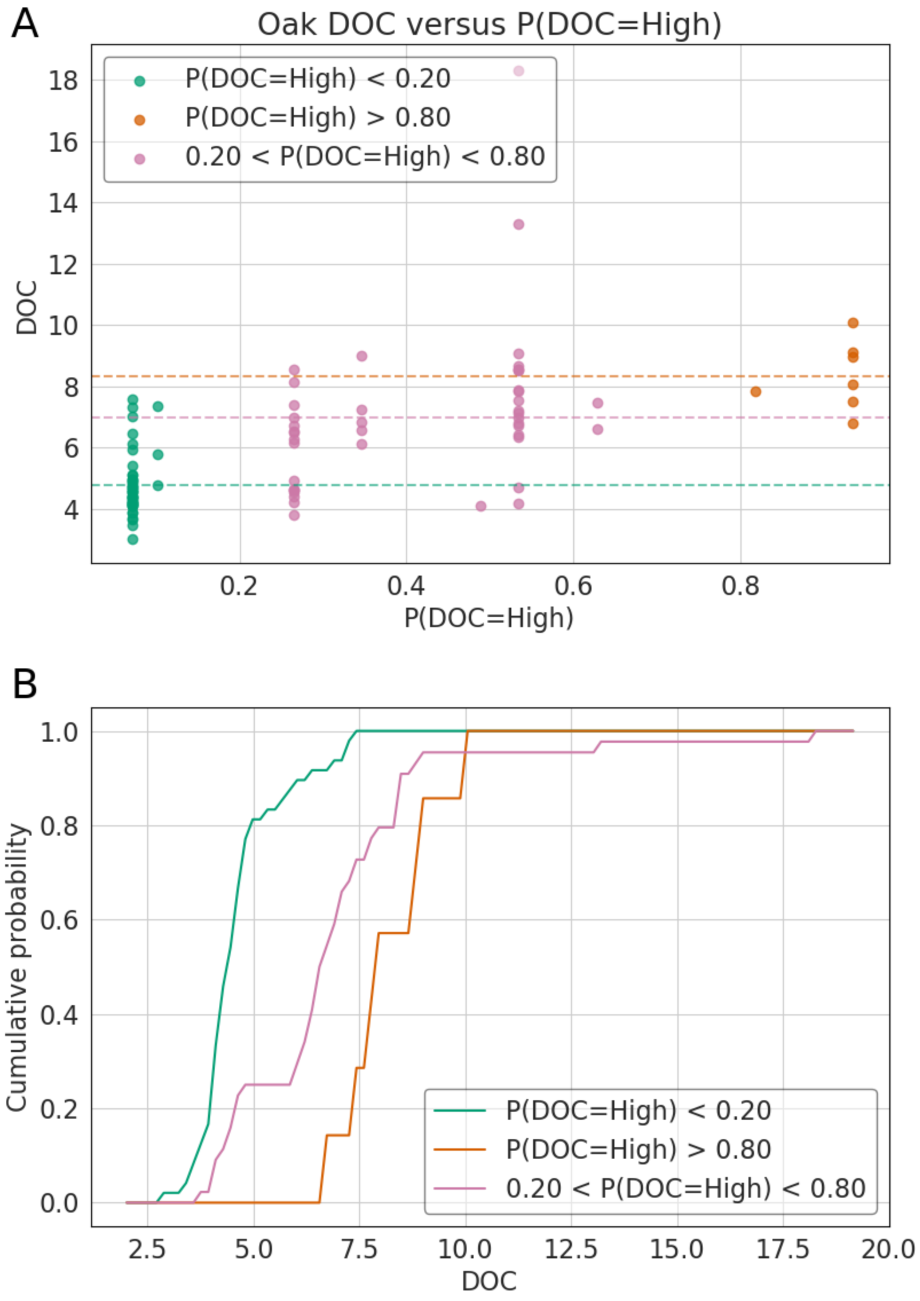


**S2 Figure. Prediction performance of 2 feature model on oak data** (A) Samples predicted with high confidence to have low DOC (P(DOC=High) $< 0.2$) and high DOC (P(DOC=High) $> 0.8$) are shown in green and orange, respectively. Predictions for which confidence is less certain are shown in magenta. Dotted horizontal lines represent the average measured DOC for each group.(B) The set of measured DOC values in the oak data set binned into low, uncertain, and high DOC categories according to the predicted probability of high DOC using a BN trained on pine samples. The Kolmogorov-Smirnov (K-S) distance between true DOC values of samples with a high predicted DOC and DOC values of samples with low predicted DOC was 0.92, p-value $= 1.69 * 10^{-5}$.

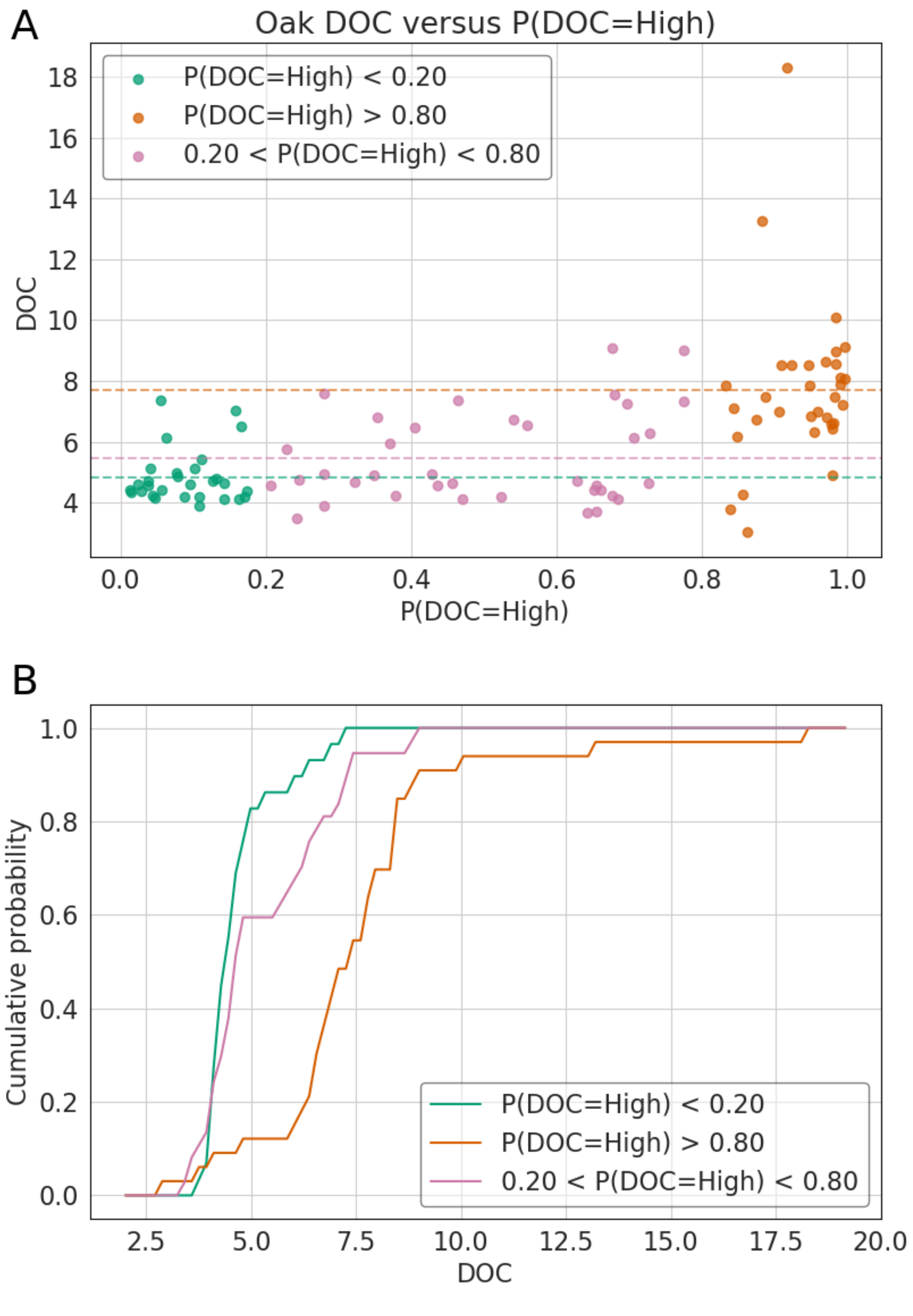


**S3 Figure. Prediction performance of 12 feature model on oak data** (A) Samples predicted with high confidence to have low DOC (P(DOC=High) $< 0.2$) and high DOC (P(DOC=High) $> 0.8$) are shown in green and orange, respectively. Predictions for which confidence is less certain are shown in magenta. Dotted horizontal lines represent the average measured DOC for each group.(B) The set of measured DOC values in the oak data set binned into low, uncertain, and high DOC categories according to the predicted probability of high DOC using a BN trained on pine samples. The Kolmogorov-Smirnov (K-S) distance between true DOC values of samples with a high predicted DOC and DOC values of samples with low predicted DOC was 0.78, p-value $4.19 * 10^{-9}$.

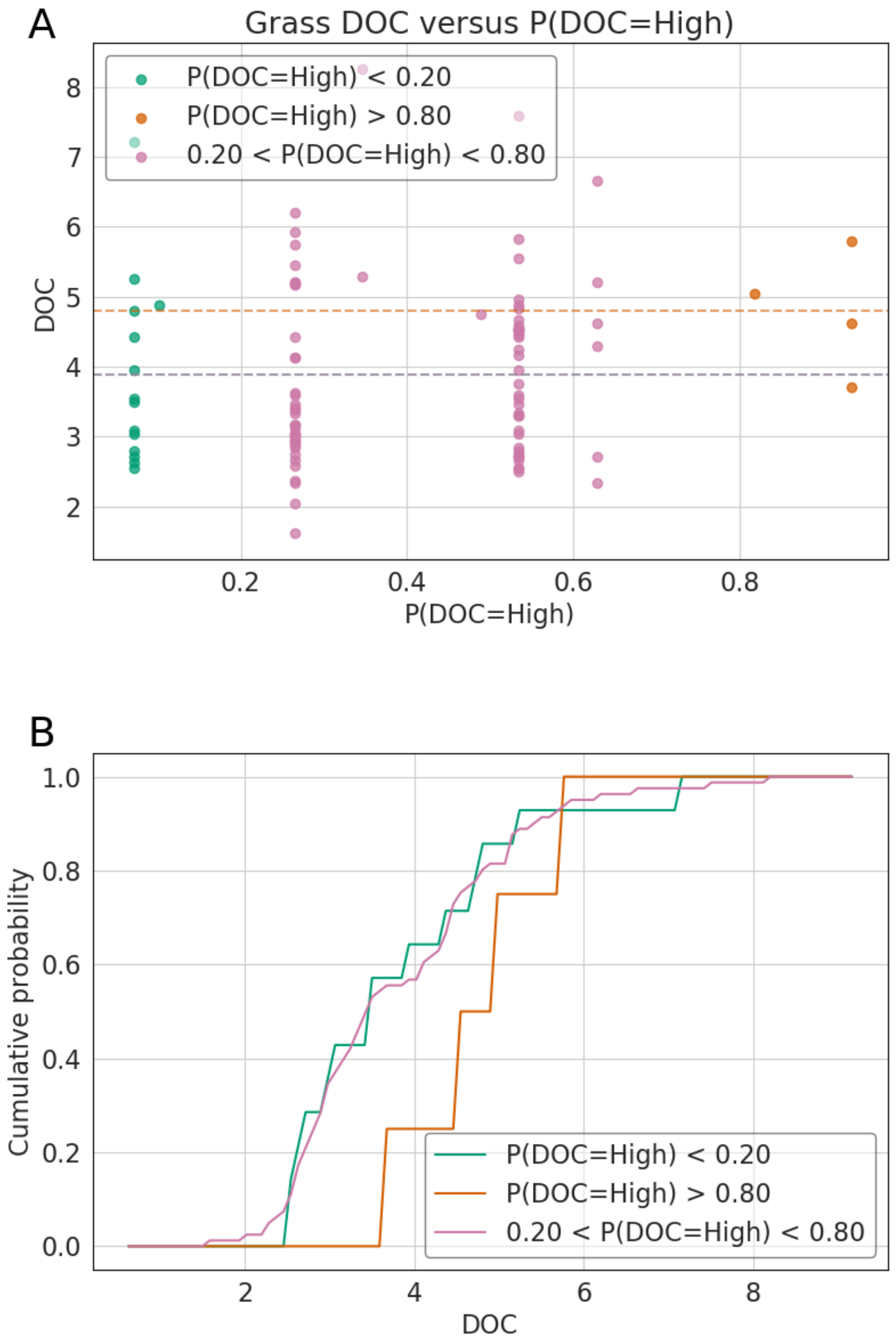


**S4 Figure. Prediction performance of 2 feature model on grass data** (A) Samples predicted with high confidence to have low DOC (P(DOC=High) $<$ 0.2) and high DOC (P(DOC=High) $>$ 0.8) are shown in green and orange, respectively. Predictions for which confidence is less certain are shown in magenta. Dotted horizontal lines represent the average measured DOC for each group.(B) The set of measured DOC values in the oak data set binned into low, uncertain, and high DOC categories according to the predicted probability of high DOC using a BN trained on pine samples. The Kolmogorov-Smirnov (K-S) distance between true DOC values of samples with a high predicted DOC and DOC values of samples with low predicted DOC was 0.57, p-value 0.168.

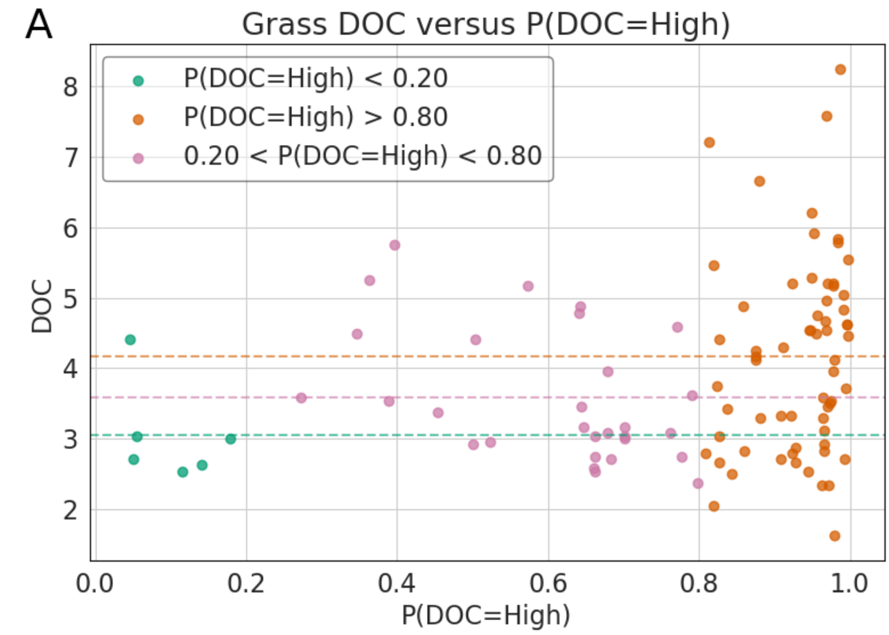


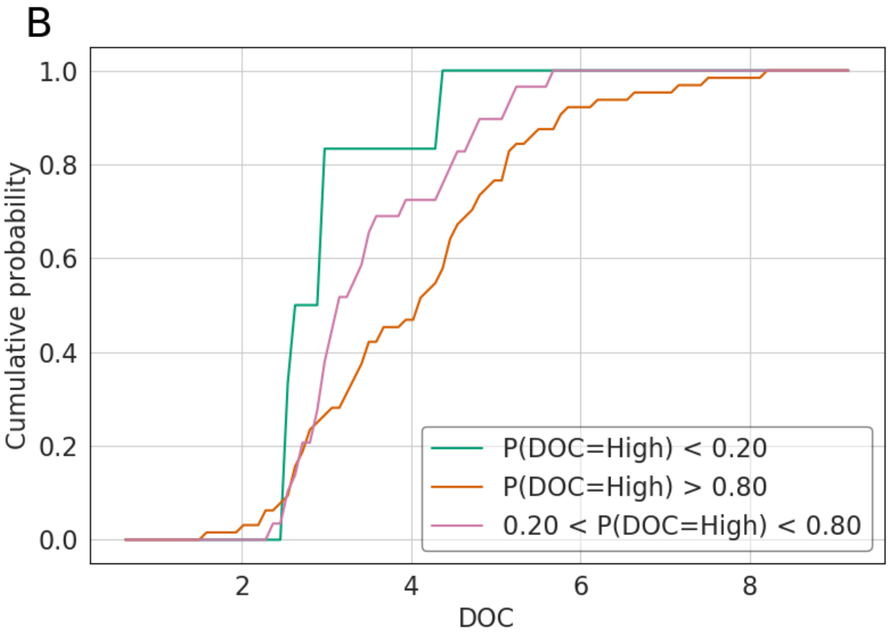


**S5 Figure. Prediction performance of 12 feature model on grass data** (A) Samples predicted with high confidence to have low DOC (P(DOC=High) $< 0.2$) and high DOC (P(DOC=High) $> 0.8$) are shown in green and orange, respectively. Predictions for which confidence is less certain are shown in magenta. Dotted horizontal lines represent the average measured DOC for each group.(B) The set of measured DOC values in the oak data set binned into low, uncertain, and high DOC categories according to the predicted probability of high DOC using a BN trained on pine samples. The Kolmogorov-Smirnov (K-S) distance between true DOC values of samples with a high predicted DOC and DOC values of samples with low predicted DOC was 0.57, p-value 0.035.

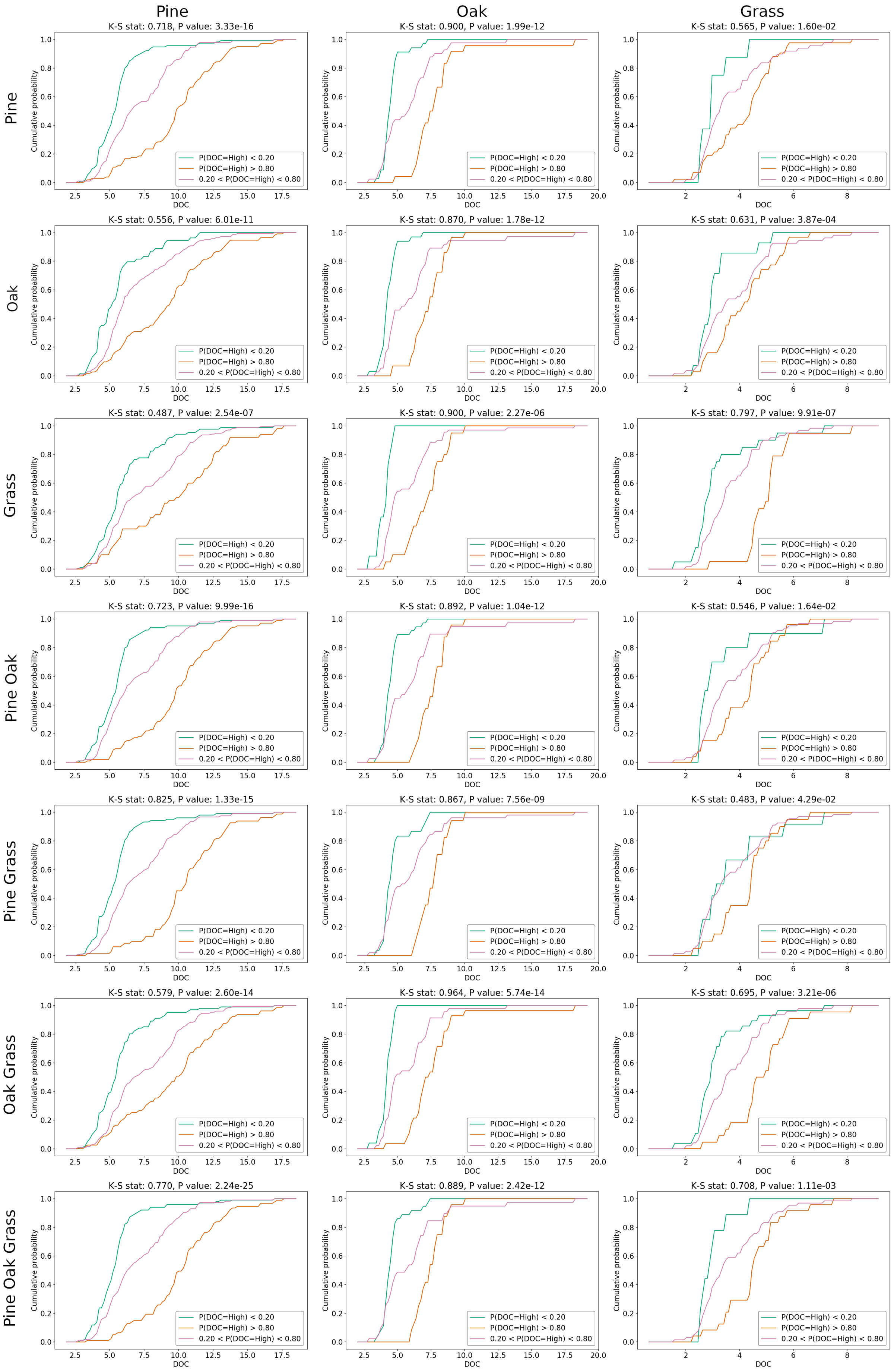


**S6 Figure. Cumulative distribution plots of out of bag predictions on pine, oak, and grass data sets using models trained on combinations of pine, oak, and grass data.** Using genera that appeared in the intersection set in pine, oak, and grass data sets, model training and prediction was performed over 50 bootstrap permutations in which ∼20% of each data set was randomly selected and withheld for testing. Each row corresponds to the training set, and each column corresponds to the testing data set.
